# Parkin overexpression attenuates muscle atrophy and improves mitochondrial bioenergetics but fails to improve key histological features in a mouse model of Duchenne Muscular Dystrophy

**DOI:** 10.1101/2025.06.13.659533

**Authors:** Olivier Reynaud, Marie-Belle Ayoub, Jean-Philippe Leduc-Gaudet, Marina Cefis, Marc P Lussier, Sabah NA Hussain, Gilles Gouspillou

**Author notes:** **Corresponding author:** Gilles Gouspillou, Département des sciences de l’activité physique Faculté des sciences, Université du Québec à Montréal (UQÀM), Pavillon des sciences biologiques (SB), Local: SB-4640 141, Avenue du Président Kennedy, Montréal, Québec, Canada, H2X 1Y4.

## Abstract

Duchenne Muscular Dystrophy (DMD) is the most common childhood muscular disorder. Mitochondrial dysfunctions are key disease features of the disease, and strategies that improve mitochondrial health have emerged as promising to slow disease progression. Emerging evidence indicates that impaired/insufficient mitophagy may contribute to the accumulation of mitochondrial dysfunction seen in patients and animal models of DMD. We therefore hypothesized that overexpressing Parkin, a key mitophagy regulator, may improve mitochondrial and muscle health in a mouse model of DMD. To this end, Parkin was overexpressed using intramuscular injections of adeno-associated viruses performed in 5-week-old and 18-week-old D2.B10-Dmd^mdx^/J mice (D2.mdx), a widely used mouse model of DMD. Four and 16 weeks of Parkin overexpression initiated in 5-week-old and 18-week-old D2.mdx, respectively, resulted in muscle hypertrophy, as indicated by an increase in muscle mass and fiber cross-sectional area. While Parkin overexpression did not impact maximal mitochondrial respiration or mitochondrial content, it increased the Acceptor Control Ratio, an index of mitochondrial bioenergetic efficiency. Parkin overexpression also decreased mitochondrial H_2_O_2_ emission, a surrogate for mitochondrial ROS production. However, Parkin overexpression failed to reduce the proportion of fibers with central nuclei and markers of muscle damage and/or necrosis. Taken all together, our results indicate that Parkin overexpression can attenuate muscle atrophy, improve mitochondrial bioenergetics and lower mitochondrial ROS production in a mouse model of DMD. These findings showcase the partial beneficial effects of overexpressing Parkin in ameliorating some, but not all, pathological features observed in a mouse model of DMD.

**Graphical abstract:** Impact of AAV-mediated Parkin overexpression on Duchenne Muscular Dystrophy (DMD) progression in skeletal muscle of D2.mdx (a mouse model of DMD). Parkin overexpression attenuated muscle atrophy, reduced mitochondrial H_2_O_2_ emissions and improved an index of mitochondrial coupling efficiency. Created with BioRender.com.

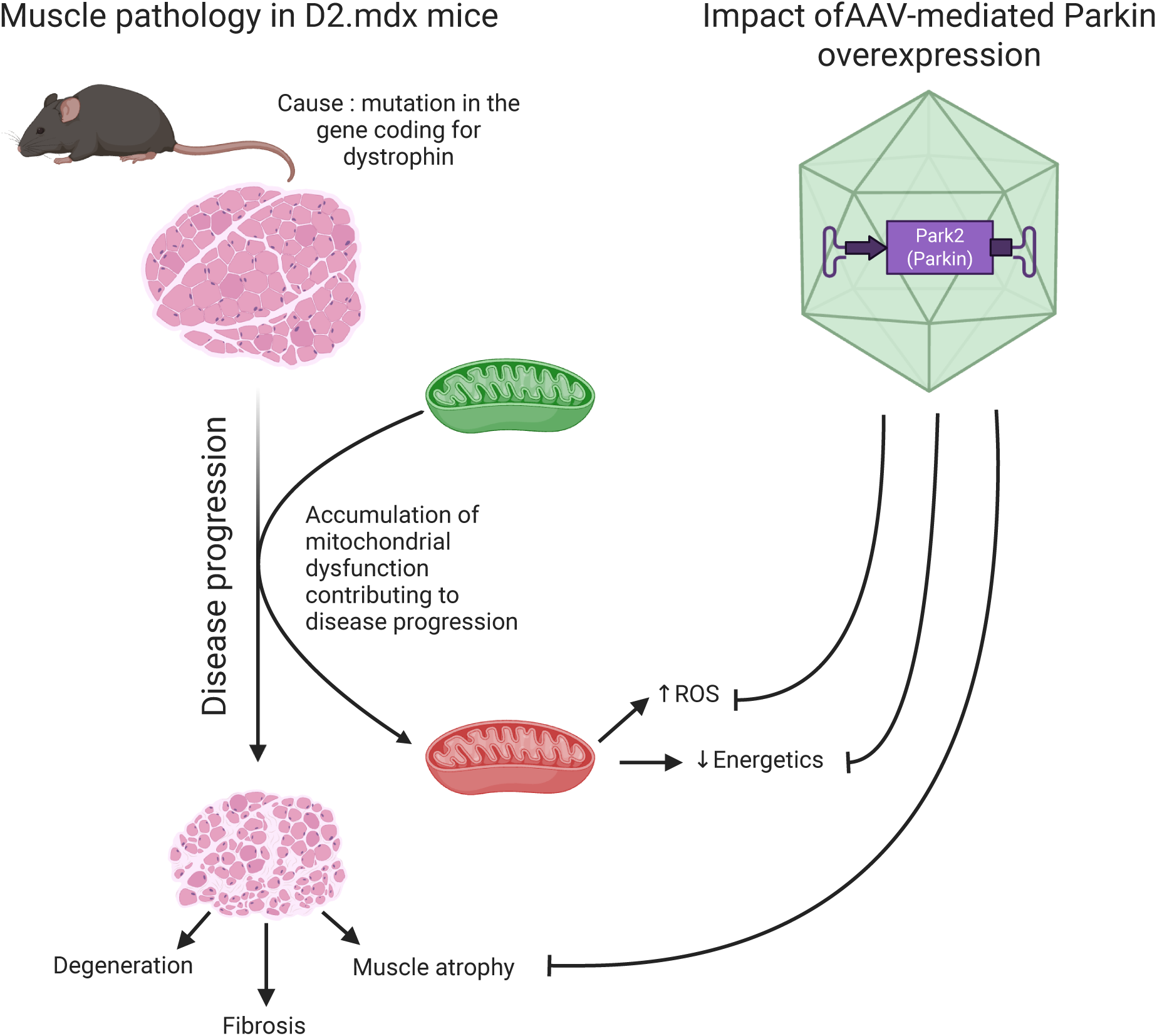

## Introduction

Duchenne muscular dystrophy (DMD) is a rare hereditary genetic disorder that affects approximately 1 in 5,000 to 1 in 6,000 live male births^1,2^ It is caused by a recessive mutation in the dystrophin gene located on the X chromosome^2,3^. DMD is characterized by progressive skeletal muscle degeneration resulting in a loss of muscular functions that can be detected around age three, and worsens over time, usually leading to loss of ambulation around puberty ^2^. The health status of DMD patients declines continuously over time until progressive loss of respiratory function and cardiomyopathy. As a result, the lifespan of DMD patients remains short, estimated to be between 22 and 28 years of age ^4^. Unfortunately, DMD currently remains an incurable genetic disease, and developing therapeutic approaches aiming at slowing disease progression and/or attenuating symptoms are regarded as useful strategies to improve the health status and quality of life of DMD patients.

Alterations in various aspects of mitochondrial function are key features observed in DMD and are thought to contribute to disease progression. Several studies have reported reduced mitochondrial content and bioenergetics^5–13^, increased ROS production ^6,7,14^, altered calcium handling ^9,10^ and increased mitochondria-mediated apoptosis ^7,10^ in mouse models of DMD. Further highlighting the key role played by mitochondrial dysfunction in DMD progression, genetic manipulations designed to improve mitochondrial health have been shown to successfully improve the dystrophic phenotype of preclinical models. Indeed, short term PGC1-α overexpression (a master regulator of mitochondrial biogenesis) by transfecting fibers of mdx mice (model of DMD) with a plasmid resulted in improved mitochondrial content, increased the capacity of mitochondria to buffer calcium, normalization of the susceptibility to permeability transition pore opening and attenuation of the proteolytic and apoptotic signalling pathways^10^. In line with these findings, mdx mice stably overexpressing PGC1-α in skeletal muscle displayed improved histological features, increased running performance and decreased plasma creatine kinase levels (a marker of muscle plasma membrane damage), while adeno-associated virus-mediated overexpression for PGC-1α in neonatal mdx mice increased muscle resistance to fatigue and to contraction-induced damage ^15,16^. Altogether, these results suggest that improving mitochondrial fitness (i.e. mitochondrial content and function) is a viable therapeutic strategy to improve muscle health in DMD.

The maintenance of mitochondrial fitness relies on the subtle coordination of processes involved in mitochondrial biogenesis (regulated in part by PGC-1α) and mitochondrial quality control processes such as mitophagy (a process responsible for the removal of dysfunctional mitochondria, regulated in part by Parkin ^17,18^). While the potential protective role of stimulating mitochondrial biogenesis has already been investigated and demonstrated ^10,15,16^, no study has investigated whether targeted increases in mitophagy regulators could also prevent the accumulation of dysfunctional mitochondria in dystrophic muscle and attenuate the progression of DMD. The need for such studies is further strengthened by recent findings indicating that mitophagy-regulating genes, including Parkin, are downregulated in dystrophic skeletal muscles^19^. We have also recently reported that Parkin overexpression can positively impact mitochondrial and muscle health in other models associated with muscle dysfunction, including aging-related and sepsis-induced muscle atrophy and weakness^20–22^. Therefore, increasing Parkin expression may represent an interesting avenue to improve skeletal muscle function in a rodent model of DMD. In this setting, we investigated the impact of Parkin overexpression, achieved using intramuscular injections of Adeno-Associated Viruses (AAVs), on skeletal muscle mass, mitochondrial respiration, and H_2_O_2_ emission in D2.B10-Dmd^mdx^/J mice (hereafter referred to as D2.mdx mice), a rodent model of DMD best mimicking the human phenotype of the pathology^23–25^. We hypothesized that Parkin overexpression would improve mitochondrial respiration, decrease ROS production, increase muscle mass, and improve histological features in D2.mdx mice.

## Materials and methods

### Animal procedures and AAV injections

The present study was carried out in strict accordance with standards established by the Canadian Council of Animal Care and the guidelines and policies of UQAM. All procedures were approved by the animal ethics committees of UQAM (#CIPA981). This study is reported in accordance with ARRIVE guidelines. Experimental protocols were designed to reduce animal suffering and number. Experiments were conducted on 5-week-old and 18-week-old male D2.mdx mice (D2.B10-Dmd^mdx^/J; JAX stock #013141), a rodent model of DMD with an early onset and pronounced dystrophy phenotype ^23,24^ or DBA/2J mice purchased from Jackson Laboratories. Two to four mice were housed per cage at 24±1°C, 50-60% relative humidity on a 12h light/dark cycle. Mice had *ad Libitum* access to a standard Chow diet and water. All AAVs used in the present study were purchased from Vector Biolabs and were of Serotype 1, a serotype with a proven tropism for skeletal muscle cells ^21,26^. After a 5-day acclimatization period, an AAV1 containing a muscle specific promoter (tMCK), a sequence coding for the reporter protein GFP and a sequence coding for Parkin (Park2) were injected intramuscularly (25 µl per site; 2.5 x 10^11^ genome copies (GC)) into the right tibialis anterior (TA) and gastrocnemius (GAS) muscles. In this construction, the sequences coding for Parkin and GFP were separated by a sequence coding for the auto-cleavable 2A peptide, allowing the separation of Parkin and GFP once translated. A control AAV1 expressing only the GFP sequence under the control of the tMCK promoter was injected in the contralateral leg. Injections were carried out under general anesthesia using 2% isoflurane. Since the AAV1 recombination site in wild-type AAV1 was deleted in these recombined AAV1s, both GFP and Parkin expressions were episomal expression without integration into the host DNA. After 4 or 16 weeks of Parkin and/or GFP overexpression, mice were anesthetized with isoflurane and subsequently euthanized by CO_2_ inhalation. The TA and GAS from both legs were removed for further experiments.

### Preparation of permeabilized muscle fibers for in situ assessment of mitochondrial function

Mitochondrial function was determined in freshly excised TA muscles as previously described ^26,27^. The muscles were dissected and rapidly immersed in ice-cold stabilizing buffer A (2.77 mM CaK_2_ ethylene glycol-bis-(2-aminoethylether)-N,N,N,N-tetraacetic acid (EGTA), 7.23 mM K_2_ EGTA, 6.56 mM MgCl_2_, 0.5 mM dithiothreitol (DTT), 50 mM 2-(N-morpholino) ethanesulfonic acid potassium salt (K-MES), 20 mM imidazol, 20 mM taurine, 5.3 mM Na_2_ ATP, and 15 mM phosphocreatine, pH 7.3) at 4°C. The muscles were weighed and then separated into small fiber bundles using fine forceps under a surgical dissecting microscope (Leica S4 E, Germany). The muscle fiber bundles were incubated into a glass scintillation vial for 30 minutes at low rocking speed containing buffer A supplemented with 0.05 mg/mL saponin to selectively permeabilize the sarcolemma. Fiber bundles were then washed 3 times 10 minutes at low rocking speed in buffer Miro 5 (0.5 mM EGTA, 6 mM MgCl2, 3 mM H20, 60 mM K-lactobionate, 20 mM Taurine, 10 mM KH2PO4, 20 mM HEPES, 110 mM Sucrose, 1 g/L fatty acid free BSA, pH 7.4 at 4°C).

### Assessment of mitochondrial respiration

The assessment of mitochondrial respiration in permeabilized TA myofiber was performed in an Oroboros O2K high-resolution fluororespirometer (Oroboros Instruments) at 37 °C in 2 mL of Miro 5 buffer as previously described ^26,27^.. Briefly, 3 to 6 mg (wet weight) of TA permeabilized fiber bundles were weighed and added to the respiration chamber. The following substrates were added sequentially: 10 mM glutamate plus 5 mM malate (G+M), 2 mM ADP, 10 mM succinate, and 400μM antimycin A. All respiration experiments were analyzed with MitoFun (https://zenodo.org/records/7510439), a homemade code designed to analyze mitochondrial function data in the Igor Pro 8 software (Wavemetrics, OR, USA) ^26^.

### H_2_O_2_ emission in permeabilized muscle fibers

H_2_O_2_ emission from TA myofiber bundles was assessed using the Amplex Ultra Red-horseradish peroxidase (HRP) system as previously described ^27^. This was performed along with the respiration assessment in the Oroboros O2K high-resolution fluororespirometer (Oroboros Instruments) at 37 °C in 2 mL of buffer Miro 5 with Amplex Ultra Red (10 μM), superoxide dismutase (SOD, 5 U/ml), and horse-raddish peroxidase (HRP, 1 U/ml) at 37°C. A calibration curve was generated daily using successive additions of known H_2_O_2_ in the absence of tissue. H_2_O_2_ emission was normalized as picomoles per minute per milligram of wet weight. All H_2_O_2_ emission experiments were analyzed with MitoFun (https://zenodo.org/records/7510439), a homemade code to analyze mitochondrial function data in the Igor Pro 8 software (Wavemetrics, OR, USA) ^26^.

### Immunoblotting

20 mg of each TA was homogenized in 10 volumes of an extraction buffer (50 mM Tris base, 150 mM NaCl, 1% Triton X-100, 0.5% sodium deoxycolate, 0.1% SDS and 1x of a protease inhibitor cocktail (Sigma-Aldrich P8340); pH 8). The homogenate was centrifuged at 12,000 *rpm* for 20 min at 4°C. Protein content in the supernatant was determined using the Bradford method using BSA as standard. Aliquots of supernatant were mixed with Laemmli buffer and subsequently heated at 95°C for 5 min. Equal amounts (20 μg) of proteins were loaded onto gradient 4-15% Mini PROTEAN® TGX Stain-Free Gels (Biorad), electrophoresed by SDS-PAGE and then transferred to polyvinylidene fluoride membranes (PVDF, Biorad). A stain-free blot image was taken using the ChemiDoc Touch Imaging System for total protein measurement in each sample lane. Membranes were incubated for 1 hr at room temperature in a blocking buffer composed of Tris-buffered saline containing 5% BSA and 0.1% Tween 20 (TBS-T; pH 7.5) and probed either overnight at 4°C or for 1 hr at room temperature with the primary antibodies. The complete list of antibodies used for immunoblotting analyses is shown in Table 1. Membranes were washed (3 × 5 min) in TBS-T and subsequently incubated with appropriate HRP-conjugated secondary antibodies (Table 1) diluted in blocking buffer for 1 hr at room temperature. Protein signals were detected using enhanced chemiluminescence substrate (Clarity ™ Western ECL substrate, 1705060, Bio-Rad), imaged with the ChemiDoc Touch Imaging System (Bio-Rad).

**Tableau 1:** Antibodies used for Western blot analysis.

| **Antibody** | **Species** | **Company** | **Cat. Number** | **Dilution** |
| --- | --- | --- | --- | --- |
| **ANT** | Rabbit | Abcam | Ab220408 | 1/1000 |
| **OXPHOS** | Mouse | Abcam | ab110413 | 1/500 |
| **PARKIN** | Mouse | Abcam | ab77924 | 1/1000 |
| **TFAM** | Rabbit | Abcam | ab131607 | 1/1000 |
| **TOM20** | Rabbit | Abcam | ab186735 | 1/5000 |
| **VDAC** | Mouse | Abcam | ab14734 | 1/1000 |
| **Goat Anti-Rabbit**<br>**IgG H&L (HRP)** | Goat | Abcam | ab6721 | 1/5000 |
| **Rabbit anti-mouse**<br>**IgG H&L (HRP)** | Rabbit | Abcam | ab6728 | 1/5000 |

### Muscle histology

Eight micron-thick cross-sections were cut in a cryostat at -20 °C and mounted on lysine-coated slides (Superfrost) to determine muscle fiber size, centrally nucleated myonuclei, as well as the proportion of necrosis areas using staining and immunohistological procedures ^27–29^.

### *In situ* determination of fiber size and localization of centrally nucleated myonuclei

The in situ determination of fiber size and localization of centrally nucleated myonuclei was performed as detailed in ^26,27^. Briefly, muscle cross-sections were first allowed to reach room temperature and rehydrated with PBS (pH 7.2) before being incubated for 15 min in a permeabilization solution (0.1% Triton X-100 in PBS). Slides were then washed three times with PBS before being incubated for 1 h at room temperature in a blocking solution with goat serum (10% in PBS). Sections were then incubated for 1 h at room temperature with rabbit IgG polyclonal anti-laminin antibody (Sigma-Aldrich L9393, 1:750). Muscle cross-sections were then washed 3 times in PBS before being incubated for 1 h with the following secondary antibody cocktail: Alexa Fluor 594 goat anti-rabbit IgG antibody (Invitrogen, A-11037, 1∶100). Sections were washed 3 times in PBS at 4°C before a 10-min incubation in a PBS solution containing 4, 6-diamidino-2-phenylindole (DAPI) (Thermo Fisher Scientific, D1306) at 4°C. Slides were then washed 3 times in PBS and finally coverslipped using Prolong Diamond (P36961; ThermoFisher Scientific) as mounting medium. Slides were imaged using an Olympus IX83 Ultra Sonic fluorescence microscope. To assess fiber size, the minimum Feret of muscle fibers was quantified by manually tracing myofibers using ImageJ (NIH, USA). An average of 388 ± 149 fibers were manually traced per muscle (minimum of fibers analyzed per muscle: 180). The proportion of fibers with at least one centrally located myonucleus was manually quantified using ImageJ.

### *In situ* determination of muscle area with damage and/or necrosis

Muscle cross-sections were stained with hematoxylin and eosin (H&E) as described in ^26^. Briefly, muscle cross-sections were first allowed to reach room temperature, washed with 95% ethyl alcohol, and then fixed with formalin (10% in PBS). Sections were then washed with distilled water and then incubated in hematoxylin stain (Harris Modified) for 30 sec. Sections were again washed with distilled water and 95% ethyl alcohol and then incubated in eosin Y for 15 sec. After rinsing with 95% ethyl alcohol followed by 100% ethyl alcohol, sections were cover-slipped with aqueous mounting medium. Slides were imaged with a Zeiss fluorescence microscope (Zeiss Axio Imager 2). The proportion of areas with signs of damage and/or necrosis was manually quantified using ImageJ.

### Statistical analysis

All statistical analyses were performed using GraphPad Prism 10.1.2 (GraphPad Software Inc., La Jolla, CA, USA). When only one variable was compared between muscles expressing GFP vs Parkin in D2.mdx mice, differences were tested using paired bilateral Student’s t-tests. When only one variable was compared between control vs D2.mdx mice, differences were tested using unpaired bilateral Student’s t-tests. When multiple variables were compared, differences were analysed using standard (for comparisons involving control vs D2.mdx mice) or repeated measures (for comparisons involving muscles expressing GFP vs Parkin in D2.mdx mice) two-way analysis of variance (ANOVA) if there were no missing values or mixed-effects analyses if there were missing values. Corrections for multiple comparisons following ANOVA or mixed-effects analyses were performed by controlling for the false discovery rate using the two-stage step-up method of Benjamini, Krieger and Yekutieli. P-values <0.05 and q-values <0.05 were considered statistically significant. The exact number of animals within each group is indicated in all figure legends. All data in bar graphs are presented as means ± SEM.

## Results

### The impact of long-term Parkin overexpression on skeletal muscle mass and mitochondrial bioenergetics in adult D2.mdx mice

We first assessed whether Parkin overexpression could exert beneficial effects in skeletal muscles of adult D2.mdx mice. Parkin was overexpressed for 16 weeks using intramuscular injections of AAV in 18-week-old D2.mdx mice, an age at which muscle atrophy accelerates in these mice (Figure 1A) ^23–25^. Each animal served as its own control, with the AAV designed to overexpress Parkin injected in one leg, while a control AAV expressing GFP was injected in the contralateral leg. Parkin protein levels increased substantially in muscles injected with AAV-Parkin (Figure 1B). Parkin overexpression increased muscle mass, particularly in the GAS (Figure 1C).

**Figure 1:**
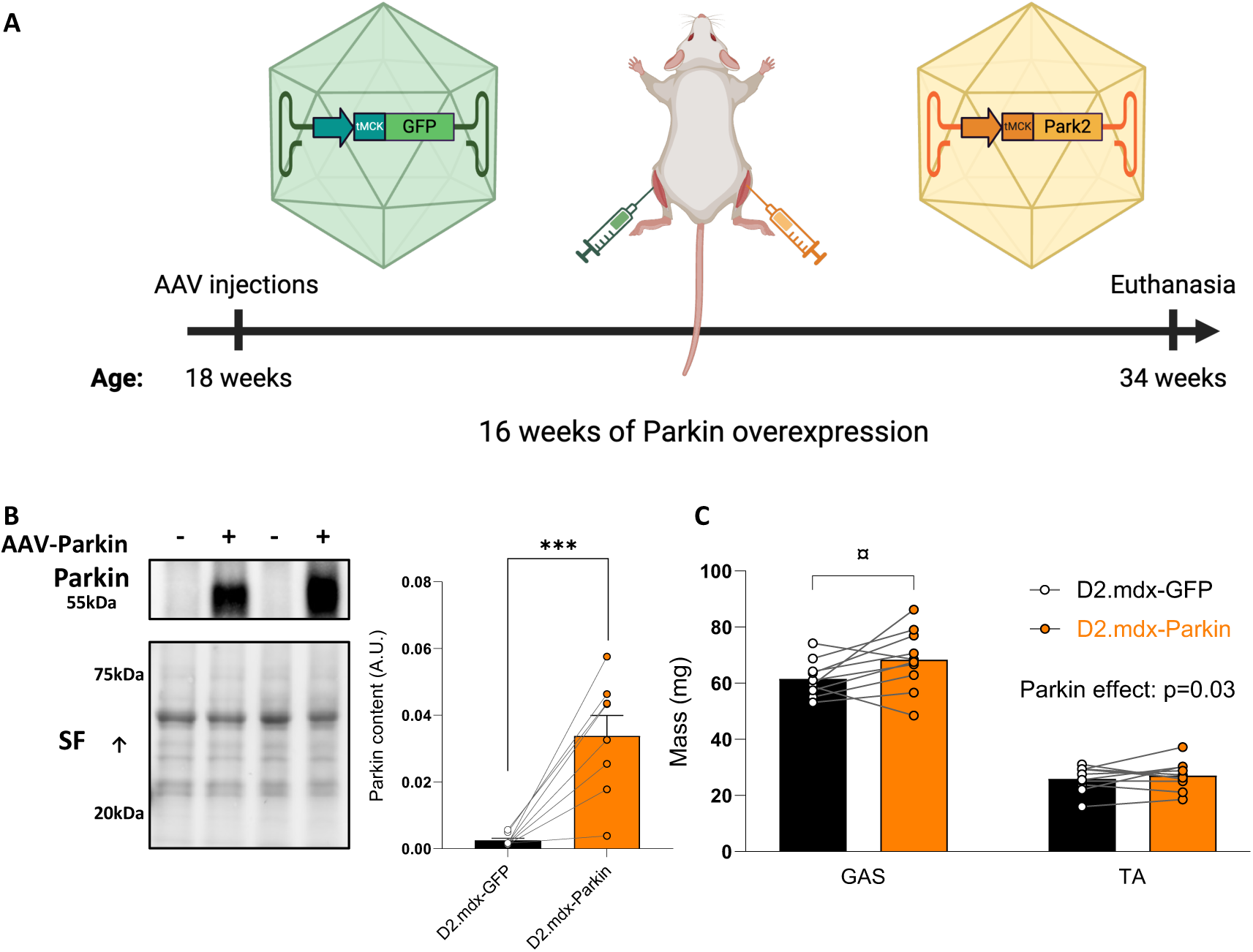
Long-term Parkin overexpression in adult D2.mdx mice increases skeletal muscle mass. **A**: Overview of the experimental design. Created with BioRender.com. **B**: Representative western blot and quantification of Parkin content in gastrocnemius (GAS) injected with AAV-Parkin or AAV-GFP of adult D2.mdx mice (n = 8/group). The stain-free (SF) technology was used to normalize protein contents. **C**: GAS and TA muscle mass injected with AAV-Parkin or AAV-GFP of adult D2.mdx mice (n = 10/group). ***p < 0.0001 (paired t-test). Q values provided in the above graphs were retrieved from two-way ANOVAs. ¤q < 0.05 (False discovery rate).

DMD is linked to reduced mitochondrial respiration ^5–7^. Because of the well-established role of Parkin in regulating mitochondrial quality control, and given that previous reports have indicated that Parkin overexpression can improve mitochondrial function in various models ^18,20–22,30^, we next investigated whether Parkin overexpression could enhance mitochondrial respiration and attenuate ROS production in muscles of D2.mdx mice. Parkin overexpression did not significantly alter basal (state II, ADP-restricted) or maximal (State III, ADP-stimulated) mitochondrial respiration assessed in permeabilized myofibers (Figure 2A). However, Parkin overexpression increased the acceptor control ratio (ACR, state III/state II respiration), a widely used index of mitochondrial coupling efficiency (Figure 2B). Since DMD has been associated with increased ROS production^6,7,14^, we also assessed mitochondrial H_2_O_2_ emission in permeabilized fibers of D2.mdx mice. Parkin overexpression decreased mitochondrial H_2_O_2_ production under the most physiologically relevant experimental conditions tested (state II and state III) (Figure 2C), but did not impact maximal (Antimycin A-induced) H_2_O_2_ emission (Figure 2D). To further investigate the impact of Parkin overexpression on mitochondrial biology in skeletal muscle D2.mdx, we next quantified various representative subunits of the oxidative phosphorylation, proteins of the outer mitochondrial membrane (ANT and VDAC1) and TFAM, a protein regulating mitochondrial DNA transcription and replication. Parkin overexpression increased the protein levels of subunits of the oxidative phosphorylation (Figure 2E). In contrast, Parkin overexpression had no impact on the content of ANT, VDAC1 or TFAM (Figure 2F).

**Figure 2:**
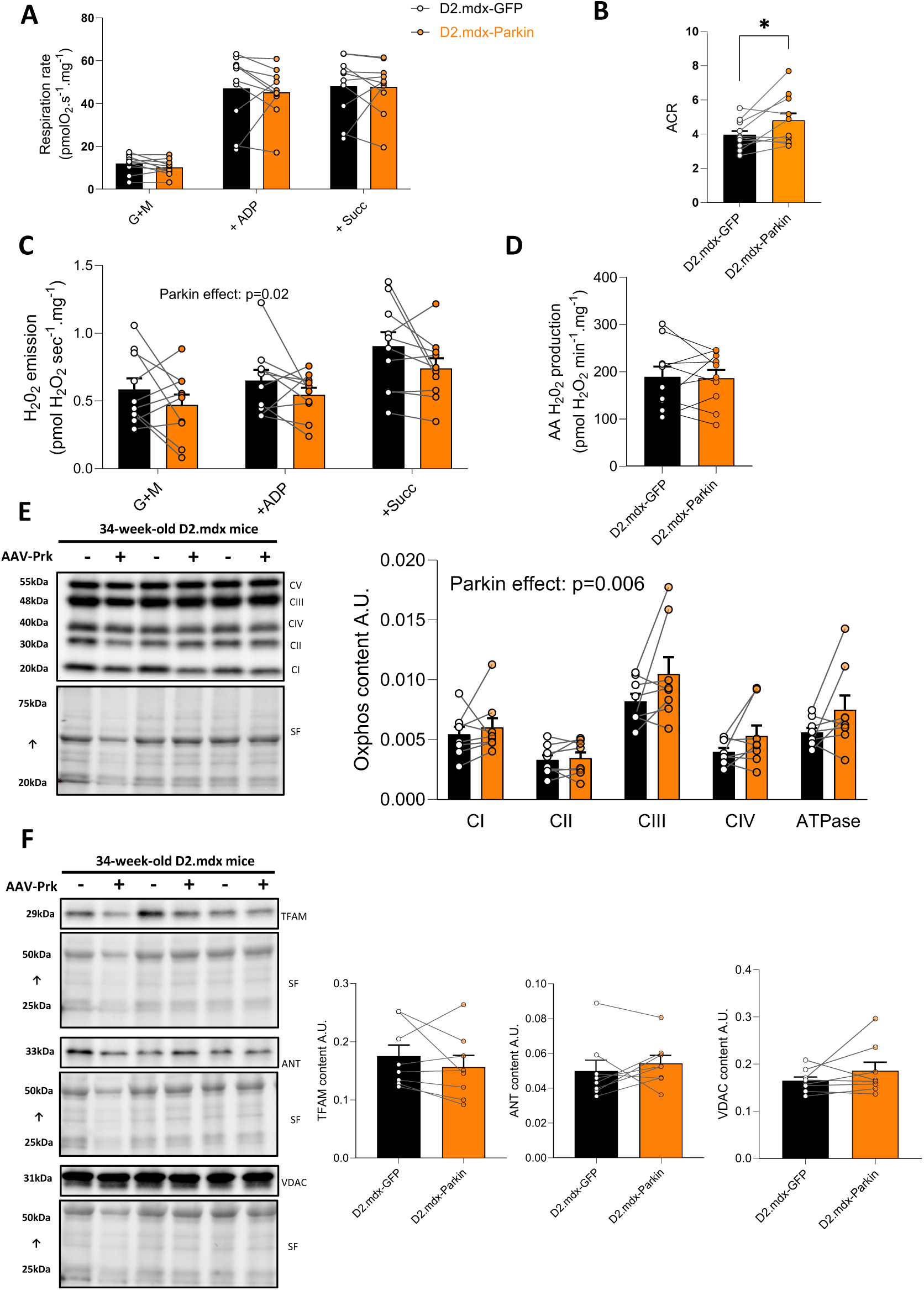
Long-term Parkin overexpression improves mitochondrial bioenergetics in adult D2.mdx mice. **A**: Quantification of mitochondrial respiration in permeabilized myofibers of the TA of D2.mdx mice using the sequential addition of Glutamate + Malate (G+M), adenosine diphosphate (ADP) and Succinate (Succ) (n = 10/group). **B**: Acceptor Control Ratio (ACR), an index of mitochondrial coupling efficiency (n = 10/group). **C**: Quantification of H_2_O_2_ emission in permeabilized myofibers driven by the sequential addition of Glutamate + Malate (G+M), adenosine diphosphate (ADP) and Succinate (Succ). **D**: Maximal (antimycin A (AA)- induced) H_2_O_2_ emission assessed in permeabilized myofibers (n = 10/group). **E:** Representative western blot and quantification of representative subunits of complexes of the oxidative phosphorylation in gastrocnemius (GAS) injected with AAV-Parkin or AAV-GFP (of adult D2.mdx mice (n = 8/group). **F**: Representative western blot and quantification of TFAM, ANT and VDAC content in gastrocnemius (GAS) injected with AAV-Parkin or AAV-GFP of adult D2.mdx mice (n = 8/group). The stain-free (SF) technology was used to normalize protein contents. *p < 0.05 (paired t-test).

These results indicate that long-term Parkin overexpression in skeletal muscles of adult D2.mdx mice increased muscle mass, improved mitochondrial bioenergetics, and lowered mitochondrial H_2_O_2_ emission.

### The impact of short-term Parkin overexpression on skeletal muscle mass, mitochondrial content and histological features in young D2.mdx mice

Based on the positive impact of Parkin overexpression in skeletal muscle of adult D2.mdx mice detailed above, we next investigated whether overexpressing Parkin at an earlier stage of the disease may confer greater protection. Parkin was overexpressed for 4 weeks in 5-week-old D2.mdx (Figure 3A), an age period during which degeneration, regeneration and inflammation are at their highest and fibrosis is progressively increasing in D2.mdx mice ^23,25^. Aged-matched DBA/2J mice (the wild type littermates of D2.mdx, thereafter referred to as control mice) injected with control AAV (AAV-GFP) were also included for comparisons. Young D2.mdx mice displayed clear signs of muscle atrophy in the GAS and TA (Figure 3C&D). Intramuscular injection of AAV-Parkin successfully and robustly increased Parkin content in young D2.mdx mice (Figure 3B). Parkin overexpression also resulted in a significant increase in the mass of the GAS (Figure 3C&D) but not in the TA (Figure 3E&F).

**Figure 3:**
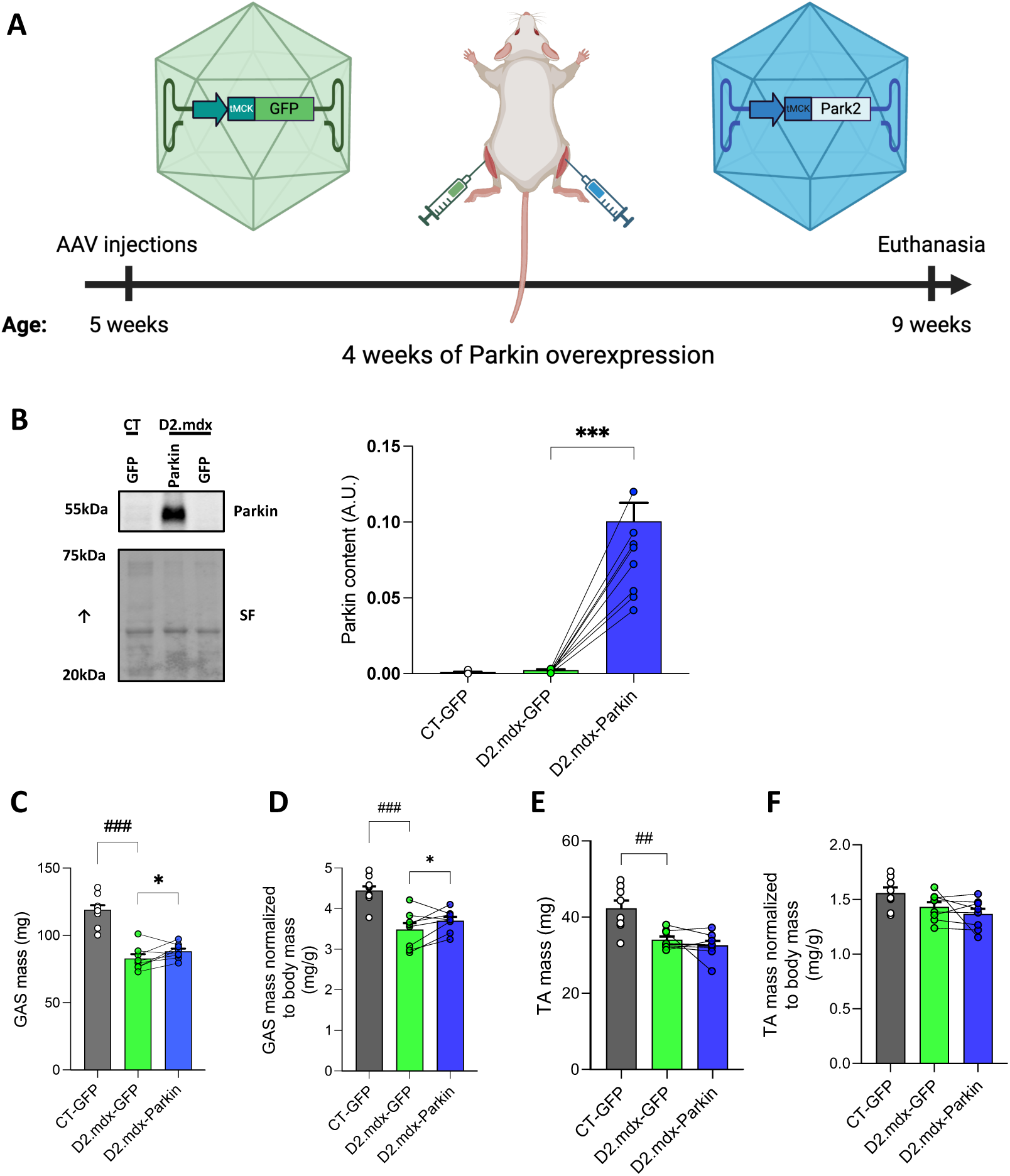
Short-term Parkin overexpression increases muscle mass in young D2.mdx mice. **A**: Overview of the experimental design. Created with BioRender.com**. B**: Representative western blot and quantification of Parkin content in gastrocnemius (GAS) injected with AAV-Parkin or AAV-GFP of young D2.mdx (n = 8/group) and control (CT; n = 10) mice. The stain-free (SF) technology was used to normalize protein contents. **C**: Absolute mass (Left) and **D**: Mass normalized to body weight (Right) of GAS injected with AAV-Parkin or AAV-GFP of young D2.mdx (n = 8/group) and CT (n=10) mice. **E**: Absolute mass (Left) and D: Mass normalized to body weight (Right) of TA injected with AAV-Parkin or AAV-GFP of young D2.mdx (n = 8/group) and CT (n = 10) mice. *p < 0.05, ***p < 0.0001 (paired t-tests). ###p<0.0001, ##p<0.001 (unpaired t-test).

However, histological analyses of the TA revealed that Parkin overexpression increased the average myofiber size in D2.mdx, indicative of attenuated muscle atrophy in Parkin overexpressing TAs of D2.mdx mice (Figure 4A-B). Similarly, upon overexpression of Parkin in D2.mdx skeletal muscle, a shift to the right in myofiber distribution was observed. in TA of D2.mdx (Figure 4C). However, it is interesting to note that while Parkin overexpression increased myofiber size in D2.mdx mice, it did not rescue the altered shape of the fiber size distribution characteristic of dystrophic muscles (i.e. a fiber type distribution characterized by a high proportion of small and large fibers; Figure 4C).

**Figure 4:**
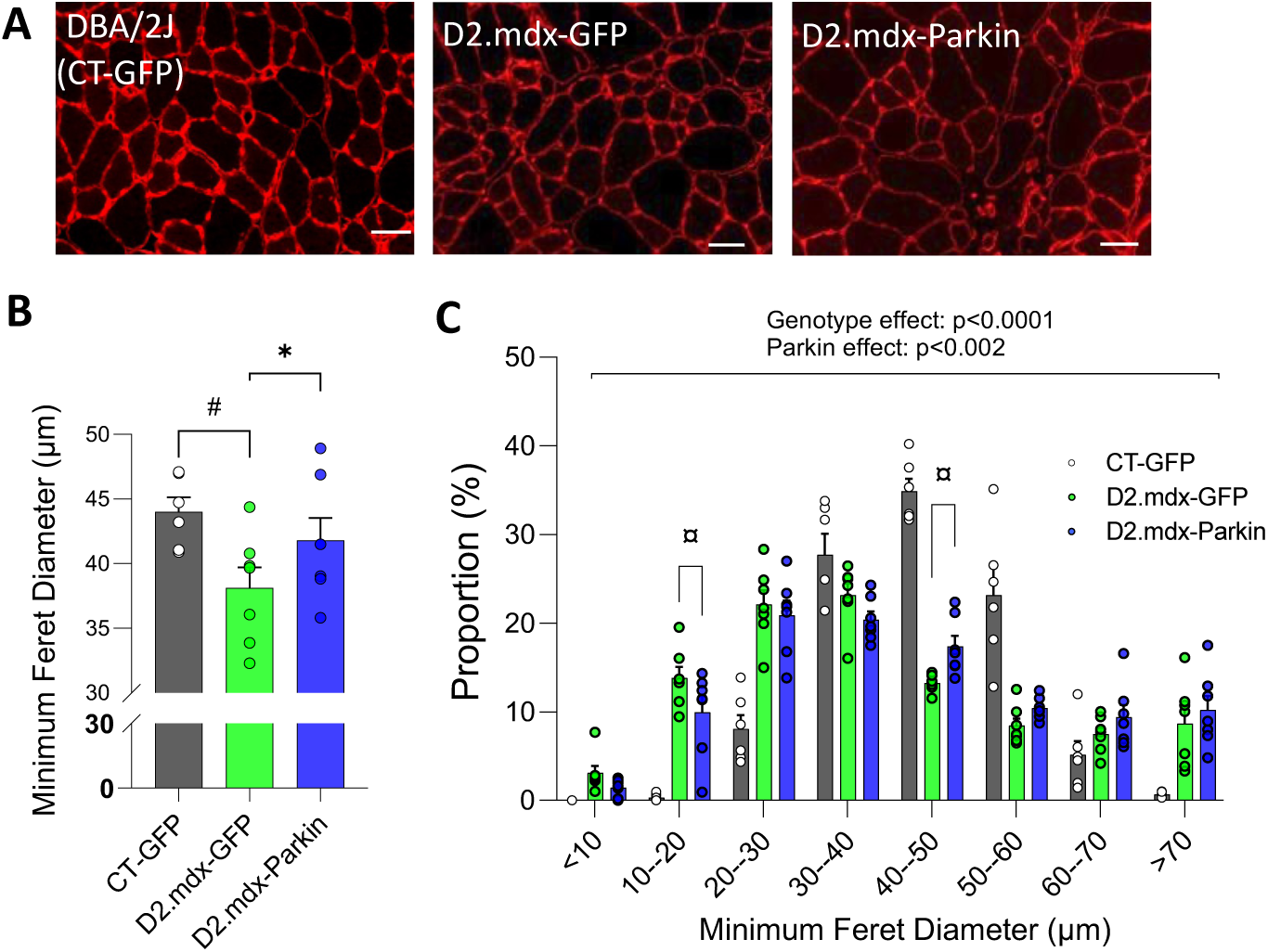
Short-term Parkin overexpression increases muscle fiber size in young D2.mdx mice. **A:** Representative laminin immunolabelling (scale bar=50µm); **B and C**: Quantification of the average minimum Feret diameter (B) and minimum Feret diameter distribution (C) for myofibers in tibialis anterior (TA) muscles injected with adeno-associated viruses (AAVs) to overexpress Parkin or GFP in control mice (n=6) or young D2.mdx (n = 7/group) mice. #p<0.05 (unpaired t-test); P values provided above graphs were retrieved from two-way ANOVAs. ¤q < 0.05 (False discovery rate).

To assess whether short-term Parkin overexpression altered mitochondrial content, we next quantified two markers of mitochondrial content: TOM20 and VDAC1 ^31^. Surprisingly, both mitochondrial markers were increased in young D2.mdx mice vs control mice (Figure 5A&B). No significant impact of Parkin overexpression on TOM20 and VDAC1 could be observed in young D2.mdx mice (Figure 5A-B).

**Figure 5:**
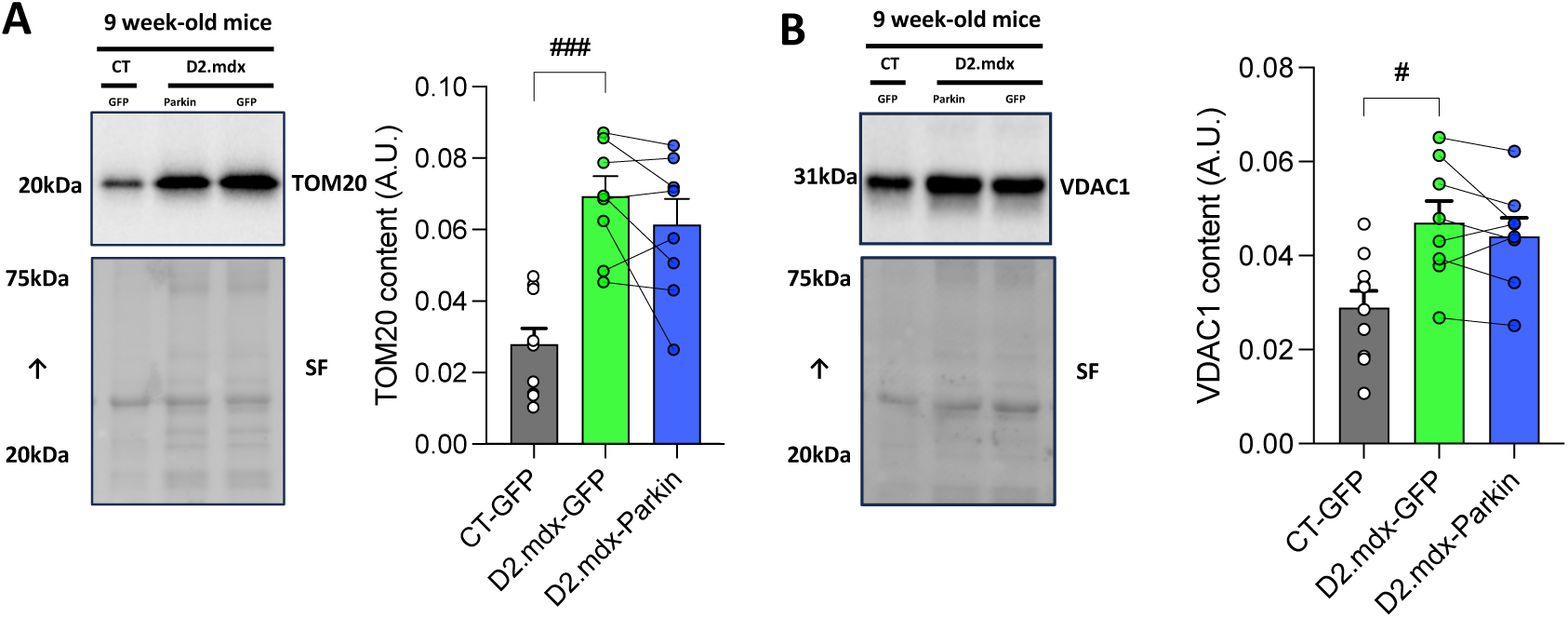
Short-term Parkin overexpression does not increase mitochondrial content in young D2.mdx mice. **A-B**: Representative western blot and quantification of TOM20 (A) and VDAC (B) content in gastrocnemius (GAS) muscles injected with AAV-Parkin or AAV-GFP of young D2.mdx (n = 8/group) and control (n = 10) mice. The stain-free (SF) technology was used to normalize protein contents. #p<0.05, ###p<0.001 (unpaired t-test).

Building on findings indicating blunted muscle atrophy in Parkin overexpressing muscle Figure 4), we next evaluated whether Parkin overexpression would reduce signs of degeneration/regeneration on histological images. Parkin overexpression did not lower the proportion of fibers with central nuclei or histological signs of damage and/or necrosis (Figure 6). Taken altogether, these findings indicate that Parkin overexpression can partly attenuate muscle atrophy in young D2.mdx but does not impact markers of mitochondrial content or classical histological features of DMD.

**Figure 6:**
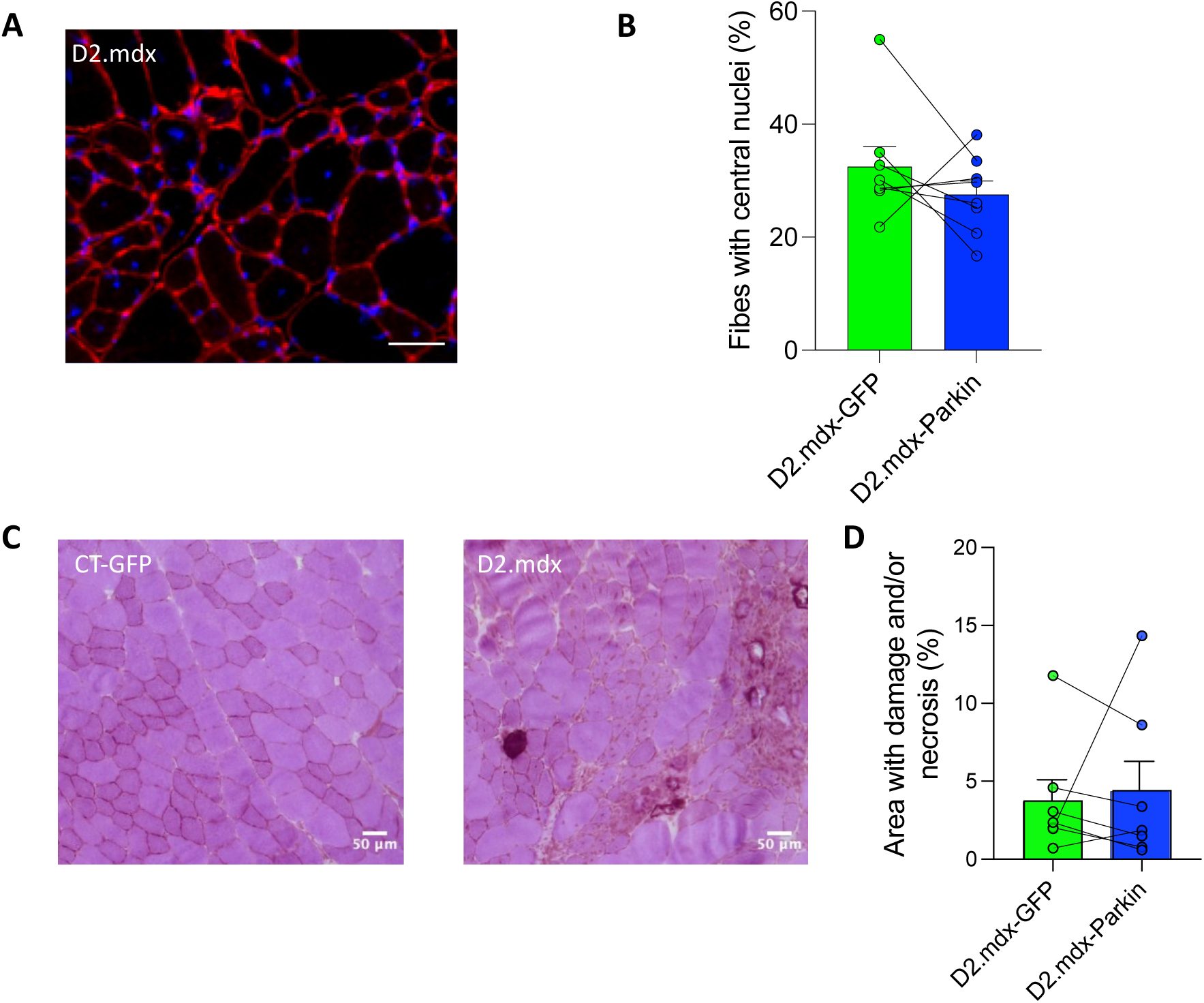
Short-term Parkin overexpression has no impact on markers of damage and/or necrosis in young D2.mdx mice. **A**: Representative image showing DAPI staining (blue) and laminin immunolabelling (red) of muscle cross sections in young D2.mdx mice. **B**: Quantification of the percentage of fibers with central nuclei in the Tibialis Anterior (TA) muscles injected with adeno-associated viruses (AAVs) to overexpress Parkin (D2.mdx-Parkin) or GFP (D2.mdx-GFP) in young D2.mdx mice (n = 8/group). **C:** Representative images of Hematoxylin and Eosin staining from muscle cross-sections in young D2.mdx mice. **D**: Corresponding quantification of the percentage of necrotic area in the TA muscles injected with AAVs to overexpress Parkin (D2.mdx-Parkin) or GFP (D2.mdx -GFP) in young D2.mdx mice (n = 7/group).

## Discussion

Alterations in multiple aspects of mitochondrial function are key characteristics of DMD, thereby positioning strategies that can improve mitochondrial health as a promising research avenue to slow disease progression ^2,32,33^. This view is supported by available data showing that overexpressing PGC-1α, a key regulator of mitochondrial biogenesis, can have beneficial effects in preclinical models of DMD ^10,15,16^. However, whether overexpressing proteins involved in the elimination of damaged or dysfunctional mitochondria through mitophagy can also exert beneficial impacts in preclinical models of DMD has never been investigated. The need for such research is further highlighted by recent data showing that genes involved in mitophagy, including Parkin, are downregulated in patients with DMD and in mdx mice ^19^. In this setting, the present study therefore aimed at assessing the impacts of Parkin overexpression, achieved with intramuscular injections of AAVs in D2.mdx mice, arguably the mouse model most closely mimicking disease progression in humans ^25^. We report here that Parkin overexpression increased dmuscle mass in both young and adult D2.mdx mice. While Parkin overexpression did not alter mitochondrial content or maximal respiration, it nonetheless increased indices of mitochondrial coupling efficiency and lowered mitochondrial H_2_O_2_ emission, indicating that Parkin overexpression positively impacted mitochondrial bioenergetics. However, despite its positive impact on muscle mass and myofiber size, Parkin overexpression failed to alter key histological characteristics of DMD, manifested as an increased number of fibres with central nuclei or markers of muscle damage and/or necrosis.

Previous studies have reported that parkin overexpression can increase muscle mass and myofiber size in healthy adult mouse skeletal muscles^21,22^, attenuate the aging-related loss of muscle mass and function in mice and C.elegans ^21,30,34^ and attenuate sepsis-induced muscle atrophy ^20^. Herein, we found that Parkin overexpression leads to an increase in muscle mass in both young and adult D2.mdx mice and increases myofiber size in young D2.*mdx* mice. These results not only extend the existing literature by positioning Parkin as an important regulator of muscle health ^21,30,35–37^, but also indicate that overexpression of Parkin can partly mitigate the muscle atrophy characteristic of DMD. It should be noted, however, that the absence of data on skeletal muscle contractility is a limitation of the present study, as we were unable to establish whether Parkin overexpression is effective in increasing muscle strength in D2.mdx mice.

While we previously reported that Parkin overexpression can increase mitochondrial content and maximal mitochondrial respiration in healthy adult skeletal muscles^21,22^, no impact of Parkin overexpression on these mitochondrial parameters were observed in dystrophic skeletal muscles. However, Parkin overexpression in skeletal muscles of D2.mdx mice increased the ACR, a widely used marker of mitochondrial coupling efficiency, indicating a positive impact on mitochondrial bioenergetics. Considering that mitochondrial bioenergetics defects are key features of DM^5–10^, improving mitochondrial efficiency may prove beneficial in attenuating disease progression.

Another area where parkin overexpression exerts beneficial effects surrounds mitochondrial ROS production. Increased mitochondrial ROS production is indeed another key feature of DMD ^6,7,14^. We report herein that Parkin overexpression lowers mitochondrial H_2_O_2_ emission, a surrogate for mitochondrial ROS production, under the most physiologically relevant experimental conditions tested (states II and III) in adult D2.mdx. The impact of Parkin overexpression aligns with our previous findings, which show that Parkin overexpression reduces mitochondrial H_2_O_2_ emission in healthy skeletal muscles ^22^ and lowers markers of oxidative stress in old skeletal muscles^21^. The positive impacts of Parkin overexpression on mitochondrial bioenergetics and ROS production were accompanied by a mild but significant increase in the content of representative subunits of the oxidative phosphorylation. While it is surprising that this increase was not associated with an improvement in the maximal mitochondrial respiration, these data may suggest that Parkin overexpression exerted, at least in part, its beneficial effects by improving the turnover of subunits of the oxidative phosphorylation and consequently its intrinsic functions. Further studies are required to clarify the mechanisms underlying the beneficial impacts of Parkin overexpression in D2.mdx mice. We should emphasize that non-canonical (i.e. mitophagy-independent) functions of Parkin might have contributed to the observed phenotype ^18^.

Interestingly, some of the positive impacts of Parkin overexpression in D2.mdx mice, which we report herein, are consistent with a recent study that has shown protective effects of Urolithin A, a compound known to stimulate mitophagy and increase Parkin expression, in a mouse model of DMD (mdx mice) ^19^. Indeed, Urolithin A supplementation was shown to improve mitochondrial bioenergetics and muscle fiber size in mdx mice. However, Urolithin A displayed beneficial effects that went beyond what was observed herein with Parkin overexpression, including a reduction in the proportion of fibers with central nuclei and a decrease of markers of inflammation and necrosis^19^. Our present findings therefore suggest that while an upregulation of Parkin may have contributed to the benefits of Urolithin A, its protective impacts in dystrophic muscles likely involved Parkin-independent and potentially mitophagy-independent mechanisms. The data also indicate that while Parkin overexpression attenuates muscle atrophy and improves mitochondrial bioenergetics in D2.mdx, it failed to improve key histological features of DMD.

## Conclusion

Targeting mitochondrial health has emerged in the last few decades as a promising approach to slow disease progression and improve the quality of life of DMD patients. However, investigations designed to assess the potential of manipulating proteins regulating mitophagy are scarce. In this setting, the present study reports that overexpressing Parkin, one of the most studied regulators of mitophagy, in dystrophic skeletal muscles exerts interesting beneficial effects, including an increase in skeletal muscle mass and myofiber size, improvement in indices of mitochondrial bioenergetic efficiency and a decrease in mitochondrial ROS production. However, the beneficial effects of Parkin overexpression were only partial as it failed to decrease key histological features of the disease, including the proportion of fibers with central nuclei and markers of muscle damage and/or necrosis. These findings showcase the partial beneficial effects of overexpressing Parkin in buffering some, but not all, pathological features observed in a mouse model of DMD. While Parkin overexpression alone may not be the most effective intervention to slow disease progression in DMD, it may provide additive benefits if combined with other approaches, including interventions designed to stimulate mitochondrial biogenesis in DMD muscles.

## Data Availability Statement

The data that support the findings of this study are available from the corresponding author upon request.

